# Antisense-mediated repression of SAGA-dependent genes involves the HIR histone chaperone

**DOI:** 10.1101/2021.07.05.451174

**Authors:** Julien Soudet, Nissrine Beyrouthy, Anna Marta Pastucha, Andrea Maffioletti, Zahra Bakir, Françoise Stutz

**Affiliations:** Dept. of Cell Biology, University of Geneva, 1211 Geneva 4, Switzerland

**Author notes:** **Contact Info** Correspondence (J.S.), (F.S.).

## Abstract

Eukaryotic genomes are pervasively transcribed by RNA polymerase II (RNAPII), and transcription of long non-coding RNAs often overlaps with coding gene promoters. This might lead to coding gene repression in a process named Transcription Interference (TI). In *Saccharomyces cerevisiae* (*S. cerevisiae*), TI is mainly driven by antisense non-coding transcription and occurs through re-shaping of promoter Nucleosome-Depleted Regions (NDRs). In this study, we developed a genetic screen to identify new players involved in Antisense-Mediated Transcription Interference (AMTI). Among the candidates, we found the HIR histone chaperone complex known to be involved in *de novo* histone deposition. Using genome-wide approaches, we reveal that HIR-dependent histone deposition represses the promoters of SAGA-dependent genes *via* antisense non-coding transcription. However, while antisense transcription is enriched at promoters of SAGA-dependent genes, this feature is not sufficient to define the mode of gene regulation. We further show that the balance between HIR-dependent nucleosome incorporation and transcription factor binding at promoters directs transcription into a SAGA- or TFIID-dependent regulation. This study sheds light on a new connection between antisense non-coding transcription and the nature of coding transcription initiation.

## Introduction

Transcription and chromatin regulation are tightly linked (1). Nucleosomes, the basic units of eukaryotic chromatin, consist of a (H3-H4)_2_ tetramer flanked by two dimers of H2A-H2B, around which 147 base pairs (bp) of genomic DNA are wrapped (2). Their main role is to prevent spurious transcription by limiting access of the transcription initiation machinery to the DNA (3). Hence, eukaryotic transcription units are organized into arrays with short linkers between two nucleosomes interspersed with Nucleosome Depleted Regions (NDRs) that constitute gene promoters (1,4–6). On the other hand, transcription through chromatin is a highly disruptive process leading to partial or complete removal of nucleosomes. Some histone chaperones compensate for this disruptive process by either recycling disrupted nucleosomes or by *de novo* histone deposition (7). Hence, transcription elongation challenges chromatin integrity and consequently threatens the maintenance of a correct gene expression program.

Recent genome-wide data show that eukaryotic genomes are almost entirely transcribed by RNA Polymerase II (RNAPII) (8). Maps of nascent transcription indicate that regions previously thought to be transcriptionally silent host non-coding or read-through transcription (9–12). Thus, chromatin integrity is much more challenged than previously considered. This has two consequences. First, a large majority of nucleosomes are disrupted and/or post-translationally modified by transcriptional read-through (6). Second, the transcription of many long non-coding RNAs (lncRNAs) overlaps with functional elements such as promoters of coding genes, potentially leading to Transcription Interference (TI) *i.e*. repression of the coding gene (13).

Transcription of lncRNAs can invade coding gene promoters either *in tandem* or in antisense configurations with respect to the coding gene. High levels of nascent antisense transcription into promoters in Eukaryotes correlate with significantly lower levels of coding sense transcription (13–15). Thus, we recently proposed antisense transcription as one parameter significantly regulating 20% of the *S. cerevisiae* coding genes (16). Several molecular scenarios could explain Antisense-Mediated Transcription Interference (AMTI) such as collisions of RNA Polymerases II (RNAPII), removal of sense transcription initiation machineries by the passage of the antisense transcription machinery or specific RNA secondary structures at promoters induced by antisense transcription (17). However, one clue that drew attention is the regulation of chromatin as loss of several chromatin regulators is known to alleviate AMTI (18–22). We proposed a mechanism in which antisense extension into promoter NDRs induces nucleosome repositioning *via* the deacetylation of nucleosomes flanking the NDRs (16). The nucleosome shift leads to Pre-Initiation Complex (PIC) binding hindrance. Hence, antisense non-coding transcription controls coding sense expression, at least partly, through the re-shaping of promoter NDRs.

The steady-state architecture of eukaryotic gene promoters mainly follows two configurations that are associated with different behaviors in terms of transcription (23). The first class of promoters presents an “open” configuration in which transcription factors can access the promoter. The second class is characterized by a “closed” promoter in which nucleosomes are hindering transcription factor binding. In *S. cerevisiae*, these open/closed classes of promoters are associated with the historically named TFIID- and SAGA-dependent genes respectively (24). Despite the fact that the expression of all coding genes depends on the co-activator TFIID, 10-15% of genes are also sensitive to the depletion of the co-activator SAGA (25). This class correlates with the presence of a perfect TATA-box sequence, a reduced width of promoter NDRs, a highly dynamic chromatin at promoters and a noisy gene expression (23,26–30). Although steady-state closed promoters are reminiscent of genes with high levels of antisense into their promoters (16,31), the link between SAGA-dependent genes and antisense non-coding transcription is poorly documented.

In order to define novel pathways involved in AMTI, we performed a genetic screen using the antisense-regulated *PHO84* model gene (18). Among the candidates, all subunits of the HIR histone chaperone complex were identified. The HIR complex has been involved in replication-independent (H3-H4)_2_ incorporation into chromatin both *in vitro* and *in vivo* (32,33). Although not essential, the HIR complex appears to compensate for mutations in the essential Spt6 and FACT histone chaperones (34). Thus, the proposed molecular role of the HIR complex is to deposit soluble histones when nucleosome recycling upon transcription elongation fails (35,36). However, a transcription-independent role in histone turnover at promoters has also been proposed (35–37).

Using genome-wide approaches, we show that the HIR complex represses SAGA-dependent genes through high levels of antisense transcription into promoters at steady-state. To address the mechanistic differences between SAGA and TFIID-dependent genes, we succeeded in turning a HIR and SAGA-dependent gene into a HIR-independent gene and *vice-versa* by modulation of transcription factor levels. We propose that two important features defining SAGA-dependent genes are the level of natural antisense extending into the promoters coupled to a balance in favor of HIR-dependent nucleosome incorporation instead of transcription factor binding.

## Materials and Methods

### Yeast strains, growth conditions and expression constructs

With the exception of the SGA screen for which *Saccharomyces cerevisiae* strains used for this study were derived from the *BY* background, all the different strains are derived from *W303* and *HHY* anchor away genetic backgrounds (38). Most of the cell cultures were grown in YEPD medium (1% yeast extract, 1% peptone) supplemented with 2% glucose as carbon source with the exception of the induction of newly-synthesized H3^HA^ grown in SC -URA (Figures 3, 4 and 5) and Pho4 overexpression grown in SD-TRP (Figure 6B). All strains were grown at 30°C, except those bearing the deletion of *RRP6* that were grown at 25°C. Anchor-away of Nrd1 and Reb1 was induced by adding 1 μg/ml of rapamycin to the medium for 1 hour and 30 minutes, respectively. All strains and their corresponding figures are reported in Table S2.

*PHO4* was cloned into the 2micron plasmid pRS424 using Gibson assembly (NEB). pGAL-H3-3xHA-URA3 on a single copy plasmid is a gift from the Michel Strubin lab.

### SGA screening

The detailed synthetic genetic arrary screening procedure is described in (39,40). Fresh SGA strains were pinned on top of the query strain and left to mate on enriched YEPD plates for 1 day at 25°C. Diploids were grown on minimum synthetic medium plates containing G418 and nourseothricin (clonNAT) to select for the deletion and the modified *PHO84* gene respectively, left to grow for 2 days at 25°C and then transferred to sporulation medium at 22°C for 7 days. Finally, Clon-NAT-*pho84∆::HIS3 rrp6∆::LEU2 DEL*::Kan^r^ haploids were sequentially selected on adequate selective synthetic medium over a total duration of 7 days at 25°C before being assessed for enhanced growth on His^-^ plates.

### Flow cytometry

Flow cytometry was perfomed as described in (41).

### RNA extraction and RNA-seq

RNAs were extracted using Glass-beads and TRIzol (Invitrogen). For RT-qPCR experiments, RNAs were reverse-transcribed using SuperScriptII (Invitrogen). Primers used for reverse transcription and PCR are listed in Table S2. RNA library preparation and single-end stranded sequencing were performed at the iGE3 genomics platform of the University of Geneva. All RNA-seq experiments were performed in duplicates.

### MNase-seq

The MNase-seq experiments were performed as described in (16). Libraries were prepared using NEBnext Ultra DNA library prep kit for Illumina (NEB). Samples were paired-end sequenced at the iGE3 genomics platform of the University of Geneva.

### ChEC-seq

The experiment was performed as described in (16) except that induction of MNase activity by addition of calcium was performed for 1 min and 5 min for the Hir2-ChEC, 30 sec for the TBP-ChEC and 5 min for the free-MNase control. Samples were then sequenced using a paired-end approach at the iGE3 genomics sequencing platform of the Univerrsity of Geneva.

### Induction of newly synthesized H3HA and MNase-ChIP-seq

The Nrd1-AA strain transformed with the pGAL-H3-3xHA-URA3 plasmid (gift from Michel Strubin lab) was grown overnight in SC -URA +2% Raffinose at 30°C. Cells were then diluted to OD_600_=0.2 to resume exponential growth until OD_600_=0.4. α-factor (20ng/ml final concentration) was added to the culture for 3h at 30°C to synchronize cells in G1-phase. Galactose powder (2% final concentration) was then directly added to the medium to induce H3^HA^ synthesis and the culture was split to add or not rapamycin (1 μg/ml final) for 1h. Cells were then processed for MNase-ChIP-seq as in (16). Newly synthesized H3^HA^ was immunoprecipitated with an anti-HA antibody (anti-HA.11, 901502, BioLegend).

### ChIP-qPCR

The experiments were performed as described in (16), without addition of any spike-in. DNA fragments were amplified with the different oligos listed in Table S2 using the SYBR Green PCR Master Mix (Applied Biosystems) and a Real-Time PCR machine (Bio-Rad).

### List of genes, TBS and nucleosomes coordinates

The list of gene coordinates from TSS to poly-A was kindly provided by the Mellor Lab. Among them were picked the ones considered as “Verified” genes in the *Saccharomyces* Genome Database (SGD) giving a complete list of 4,775 coding genes. For the TBS coordinates, our list was crossed with the ChIP-exo data from the Pugh lab (29). +1 nucleosome coordinates were extracted from DANPOS2 analysis with default settings (42) and from H3K18ac profiles of our previous publication (16). Figures 3A-C were obtained using the coordinates of the nucleosome atlas from the Friedman lab (43). The list of SAGA- and TFIID-dependent genes was obtained from (24) and crossed with our list of “Verified” genes. CR and TATA/TATA-less genes were picked in (25). Reb1 regulated genes were obtained from (44) *via* analyses of RNA-seq upon Reb1 anchor-away. Genes with a 2 fold decrease were considered as Reb1 targets. Genes up-regulated upon deletion of *ASF1* were obtained from (45).

### RNA-seq analysis

Single-end reads were aligned to sacCer3 genome assembly using Bowtie2 (46) with options ‘-k 20–end-to-end–sensitive-X 800’. PCR duplicates were removed from the analysis. BigWig coverage files were generated using Bam2wig function. Differential expression analysis was performed using the R/Bioconductor package DEseq on mRNA annotations Ensembl (*Saccharomyces*_cerevisiae.EF4.65.gtf). Antisense transcripts with a fold-change of at least 2 and multiple testing adjusted p-value lower than 0.05 were considered differentially expressed and defined as inducible Antisense Genes (iAS). Calculations of fold-changes were subsequently performed using the density files (bigWig files).

### MNase-seq mapping

Paired-end reads were aligned to sacCer3 genome assembly using Bowtie2 (46). PCR duplicates were removed from the analysis. Then, deepTools 2.0 (47) was used through the bamCoverage function with size selection of fragments (120-200bp to visualize only proper nucleosomes and not “fragile nucleosomes” (48)), counting only the 3bp at the center of fragments and counts per million (cpm) normalization.

### ChEC-seq mapping

Adapters were first removed from the paired-end reads using the Trim Galore! Tool with default options from the Galaxy server (49). Paired-end reads were then aligned to sacCer3 genome assembly using (46). PCR duplicates were removed from the analysis. DeepTools 2.0 (47) was then used through the bamCoverage function with size selection of fragments (0-120bp for TBP-ChEC and mainly 120-200bp for Hir2-ChEC) and counting of only the 3bp at the center of fragments.

### Metagene analyses and Heatmaps

Metagene plots were produced using computeMatrix followed by plotProfile commands using DeepTools 2.0 (47). Heatmaps were produced with the help of Prism 8.0 (Graphpad).

### Data and Code Availability

The accession number for the data reported in this study is GEO: GSE<u>175991</u>.

### Statistical analyses and models design

All plots and statistical analyses of this work were performed using Prism 8.0 (Graphpad). All tests are nonpaired tests. Mann–Whitney *U* tests were used to extract a p-value. ns if p-value>0.05, ∗ < 0.05, ∗∗ < 0.01, ∗∗∗ < 0.001, ∗∗∗∗ < 0.0001. All models were designed using BioRender.

## Results

### A genetic screen to uncover new players involved in Antisense-Mediated Transcription Interference (AMTI)

We set-up a genetic screen using the *PHO84* model gene for which coding sense expression is regulated by antisense non-coding transcription. *PHO84* becomes repressed in the absence of Rrp6, the nuclear subunit of the exosome, as a result of antisense non-coding transcription elongation into its promoter (18,50). We took advantage of this simple phenotype by replacing the entire *PHO84* coding sequence by the *HIS3* gene marker (Figure 1A). The bait strain recapitulates the *PHO84* AMTI phenotype since antisense ncRNAs are stabilized and extended in the absence of Rrp6 and *HIS3* becomes repressed leading to a growth defect in the absence of histidine in the medium (Figures 1A and1B). This phenotype can be alleviated in the absence of Rpd3, a Histone DeACetylase (HDAC) known to be involved in AMTI (Figure 1B) (16,18,20). Using a classical synthetic genetic array (SGA) approach (39,40), the bait strain was then crossed with the yeast deletion library and triple mutants recovered after meiosis were assayed for growth on *HIS^-^* medium (Figure 1C).

**Figure 1:**
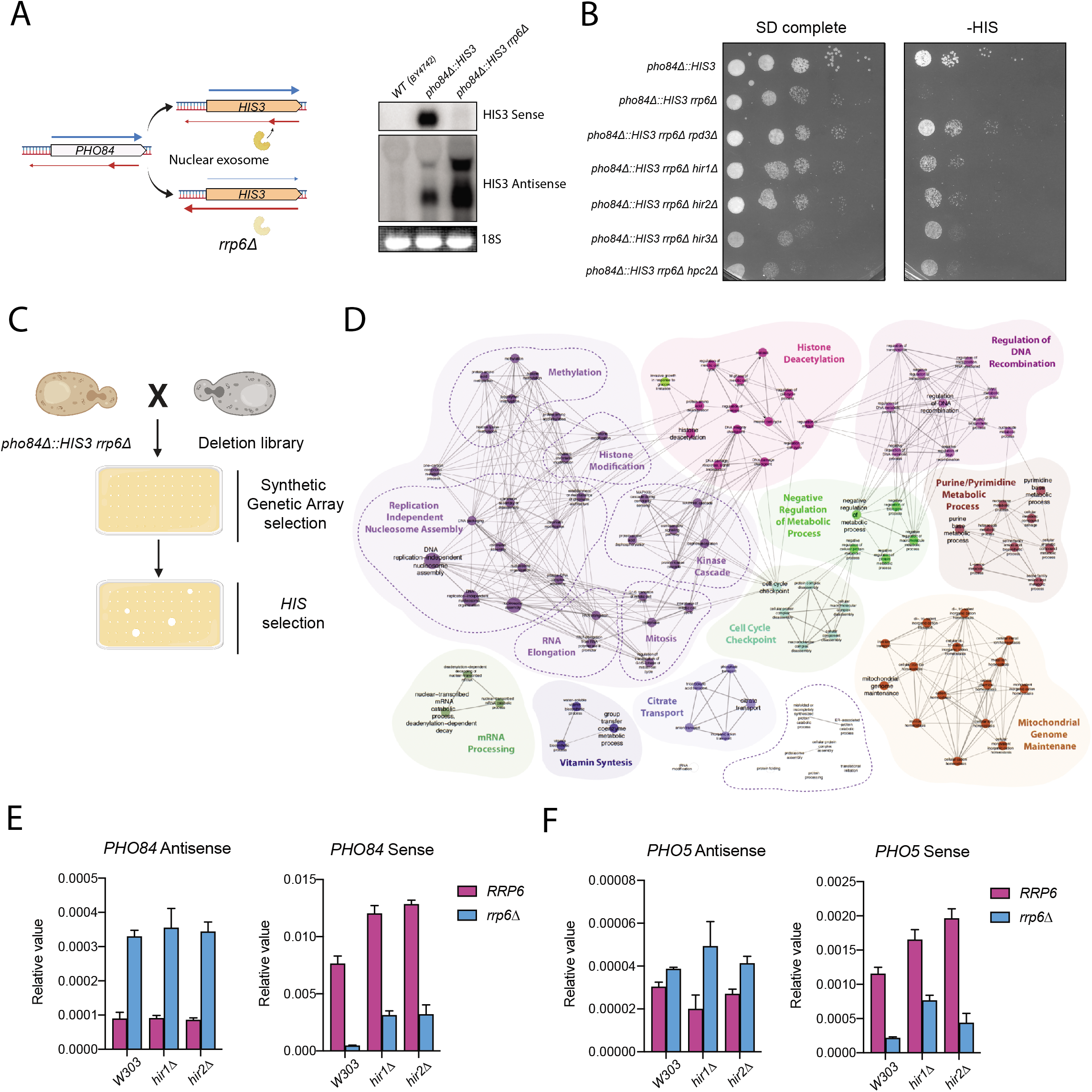
A genetic screen reveals the HIR histone chaperone complex as involved in *PHO84* and *PHO5* induced AMTI. **(A)** Design of the genetic screen and validation by Northern blotting. The *PH084* coding sequence was replaced by the *HIS3* marker. In presence of the nuclear exosome, *HIS3* antisense is early terminated and poorly triggers AMTI. When the nuclear exosome is inactivated through *RRP6* deletion, *HIS3* antisense extends into the promoter triggering AMTI. For the Northern blot, a probe targeting either *HIS3* Sense or Antisense was used. The 18S rRNA visualized by ethidium bromide staining after gel migration serves as a loading control. **(B)** Growth assay on plates. 10-fold dilutions of the indicated strains were spotted on either SD complete or HIS^-^ media and grown for 1 day at 25°C. **(C)** Synthetic Genetic Array strategy. The *pho84Δ::HIS3 rrp6Δ* strain was crossed with the Deletion Mutant Array (DMA) (see Materials and Methods). Selected diploids were sporulated before being sequentially grown on SD and HIS^-^ media. All mutants showing enhanced growth on HIS^-^ plates were considered as positive candidates. **(D)** ClueGO representation of the genetic screen positive candidates (67,68). Each node represents an enriched GO term bearing a minimum of 4 positive candidates. The size of the node reflects the statistical relevance of the GO term. The edges in a ClueGO map indicate that the connected nodes contain proteins sharing the same enriched GO terms. **(E)** RT-qPCR analyses of *PH084* sense and antisense expression normalized to *SCRI* expression in the indicated strains. n=2, error bars represent the Standard Error of the Mean (SEM). **(F)** The same as **(E)** for sense and antisense expression of *PH05*. n=2, error bars depict SEM.

Suppression of the AMTI was observed with the deletion of 198 genes. As expected, many of the hits were related to chromatin and transcription regulation (Figure 1D and Table S1). Strikingly, all the 4 subunits of the HIR histone chaperone known to be involved in replication-independent nucleosome assembly were uncovered in this screen. AMTI was indeed attenuated when deletions of *HIR1*, *HIR2*, *HIR3* or *HPC2* genes were introduced into the bait strain (Figure 1B).

We confirmed the importance of HIR in *PHO84* mRNA repression in the W303 background combined or not with the deletions of HIR and *RRP6* (Figure 1E). Loss of either Hir1 or Hir2 leads to *PHO84* up-regulation both in the presence and absence of nuclear exosome activity. Remarkably, antisense levels are not affected by loss of Hir1 or Hir2 subunits indicating that the rescue is not a consequence of non-coding transcription down-regulation. Since the sole deletion of *HIR1* or *HIR2* leads to *PHO84* up-regulation, it raises the question of a direct link between antisense induction and HIR dependency. We observed similar phenotypes with *PHO5*, another gene regulated by antisense non-coding transcription (Figure 1F).

### Induced AMTI is alleviated at SAGA-dependent genes in the absence of HIR

To address the role of the HIR complex in antisense-mediated repression (AMTI) at the genome-wide level, we performed RNA-seq of an Nrd1 anchor-away strain (Nrd1-AA), in which Nrd1 early terminated antisense non-coding transcription can be extended into gene promoters upon rapamycin addition (16,51). This leads to a transcription interference phenotype for several hundreds of genes that we assessed in the presence or absence of Hir2. This RNA-seq experiment was highly reproducible and recapitulated the AMTI of *PHO84* and *PHO5* (Figures 2A, 2B, S1A and S1B). Moreover, by analogy with the deletion of *RRP6*, the expression pattern of these two genes follows the 1^st^ criterion (*i.e.* log_2_ Ratio (*Nrd1-AA hir2Δ*-Rap/*Nrd1-AA*-Rap)>0) and the 2^nd^ criterion (*i.e.* log_2_ Ratio (*Nrd1-AA hir2Δ* +Rap/*Nrd1-AA*+Rap)>0) (Figures 2A and 2B). We then defined the set of genes for which antisense is increased by more than 2-fold upon rapamycin treatment (Figures 2C and S1C). These 601 genes were named inducible AntiSense (iAS) genes and show a significant overlap with our previous study (Figure S1D) (16). The genes showing less than 2-fold increase in antisense were named as non-inducible AntiSense (nAS) (Figures 2C and S1C). As already described, iAS tend to undergo AMTI upon rapamycin treatment (Figure 2C).

**Figure 2:**
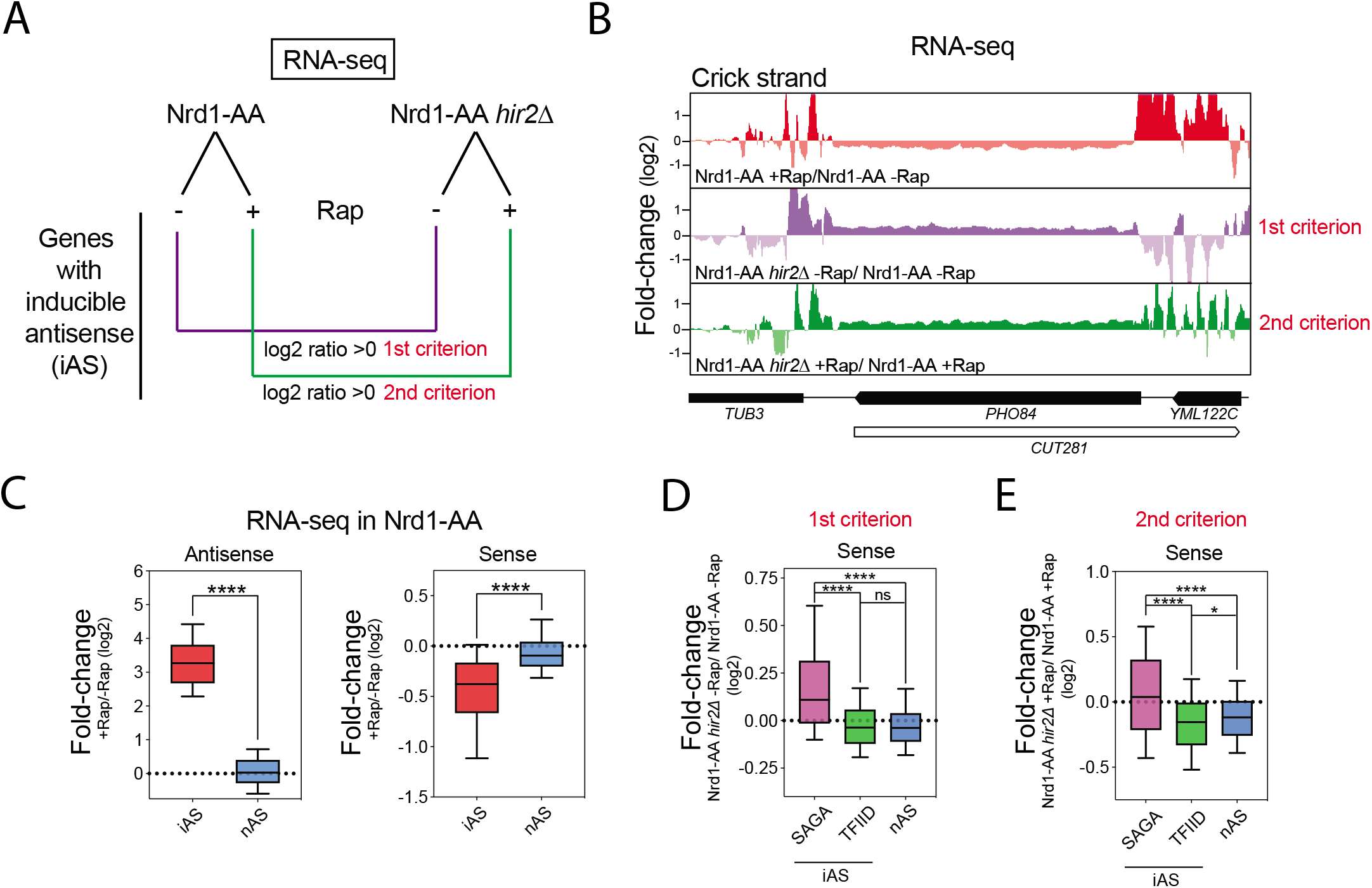
Induced AMTI is alleviated in the absence ofHIR at SAGA-dependent genes. **(A)** Design of the RNA-seq experiments using the Nrdl-AA and Nrd1-AA *hir2Δ* strains to induce AMTI at a genome-wide level. Nrdl-AA and Nrdl-AA *hir2Δ* cells were treated or not for lh with rapamycin (Rap) before RNA extraction. After definition of genes for which antisense increases upon rapamycin treatment (iAS genes), 2 criteria are used for sense expression by analogy with *PH084* and *PHOS* (**Figure 1**). 1^st^ criterion: in the absence of rapamycin, when the log_2_ fold-change Nrdl-AA *hir2Δ* /Nrdl-AA is >0. 2^nd^ criterion: in the presence of rapamycin when the log_2_ fold-change Nrdl-AA *hir2Δ* /Nrdl-*AA* is >0. **(B)** Snapshot of RNA-seq experiments depicting induced AMTI in NrdlAA +Rap/-Rap, followed by analyses according to the 1^st^ and 2^nd^ criteria at the *PH084* gene. **(C)** Boxplot showing the +Ra p/-Ra p fo ld-change of RNA-seq in the Nrdl-AA strain for the iAS genes (601 genes) and nAS genes (4,174 genes). The whole sense transcription unit was considered for calculation of fold-changes in sense and antisense. Asterisks indicate significant differences according to a two-tailed Mann-Whitney test (*P<0.05, **P<0.01, ***P<0.001, ****P<0.0001, ns: non-significant). **(D)** Boxplot showing the fold-change according to the 1^st^ criterion for the iAS SAGA-dependent genes (78 genes), iAS TFIID-dependent genes (523 genes) and nAS genes (4,174 genes). Asterisks indicate significant differences according to a two-tailed Mann-Whit ney test. **(E)** Same as in **(D)** for the 2^nd^ criterion.

Deletion of *HIR2* does not lead to AMTI alleviation following the 1^st^ and 2^nd^ criteria when iAS genes are taken as a whole (Figures S1E and S1F). However, still by analogy with *PHO84* and *PHO5*, which are SAGA-dependent genes, we split the 601 iAS genes into SAGA-dependent and TFIID-dependent classes. With this distinction, we observe a significant alleviation of AMTI for the 78 SAGA-dependent iAS genes (Figures 2D and 2E). Importantly, this phenotype cannot be attributed to a specific decrease in production of antisense in *hir2Δ* cells at SAGA-dependent genes (Figure S1G).

Thus, at a genome-wide level, the HIR histone chaperone complex contributes to induced AMTI at the iAS SAGA-dependent genes.

### Antisense induction stimulates H3 incorporation by the HIR complex

To decipher the link between antisense transcription and the HIR complex and to clarify whether they are acting in the same or parallel pathways for AMTI, we examined HIR complex binding to chromatin and its dependency on antisense induction. To do so, we performed <u>Ch</u>romatin <u>E</u>ndonuclease <u>C</u>leavage (ChEC) of the Hir2 subunit (52,53). We then paired-end sequenced the <200bp fragments protected by Hir2 and mapped them on the genome. Metagene analysis reveals a bipartite recruitment of Hir2, which is present along coding sense transcription units, following the nucleosomal pattern, but also at promoter NDRs (Figure S2A). Short induction of cleavage in ChEC experiments reveals primary binding sites while long induction indicates secondary binding sites that are released with time (52). At longer induction time points, the signal tends to accumulate at promoters indicating that the more specific binding sites correspond to the nucleosomal arrays. In order to get rid of this less specific signal, we only plotted the 120-200bp fragments, which interval contains nucleosomal particles.

We then analyzed the Hir2 ChEC-seq profile according to the levels of coding sense transcription. The Hir2 signal follows the nucleosome profile with a positive correlation between Hir2 recruitment and gene expression (Figure S2B). Since Hir2 is involved in replication independent (H3-H4)_2_ deposition on chromatin, we also analyzed the incorporation of newly synthesized H3^HA^ into chromatin of cells blocked in G1-phase and observed a nice correlation between incorporation of induced soluble H3, transcription and Hir2 recruitment (Figures S2C and S2D). Thus, HIR histone chaperone binding appears as colinear with replication-independent H3 incorporation and gene transcription.

Since Antisense induction shifts nucleosomes towards a new phasing (Figure 3A) (16), we investigated whether Hir2 recruitment is coordinately shifted upon Nrd1 anchor-away. Indeed, Hir2 relocates from its original location upon antisense induction and the same is observed for the incorporation of H3^HA^ into chromatin (Figures 3B and 3C).

**Figure 3:**
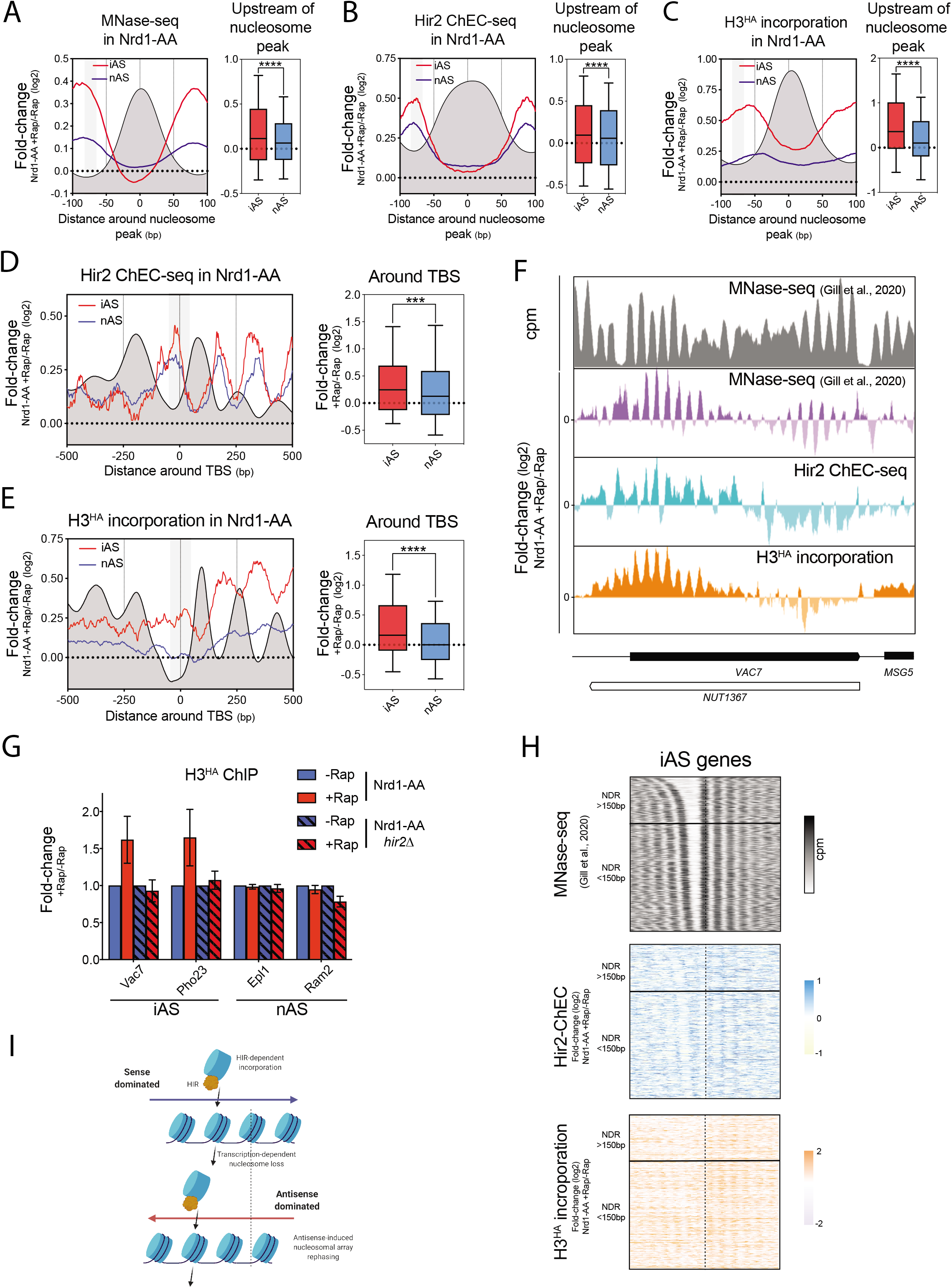
HIR binding and nucleosome incorporation follow nucleosome repositioning induced by antisense transcription. **(A)** Left: Metagene profiles of the +Rap/-Rap fold-change of MNase-seq in an Nrdl-AA strain around an average nucleosome (+1 to +6 nucleosomes are computed as an average nucleosome). Nucleosomes were averaged according to the directionality of sense coding transcription. 2,706 and 19,942 nucleosomes were considered for iAS and nAS, respectively. The dark grey profile represents the average nucleosome in the absence of rapamycin. The grey box represents the −85 to −65bp area upstream of the average nucleosome in which statistics are generated. Data were retrieved from Gill et al., 2020. Right: Boxplot showing the +Rap/-Rap fold-change for iAS and nAS in the −85 to −65bp area upstream of the average nucleosome. Asterisks indicate significant differences according to a two-tailed Mann-Whitney test. **(B)** The same as in **(A)** with Hir2 ChEC-seq data. The dark grey profile represents the Hir2 ChEC profile around the average nucleosome in the absence of rapamycin. **(C)** The same as in **(A)** with MNase-ChIP-seq data of the incorporation of newly-synthesized H3^HA^ For this experiment, ectopic H3^HA^ expression was induced for lh by galactose addition together with or without addition of rapamycin in the medium (see Materials and Methods). The dark grey profile represents the H3^HA^ incorporation around the average nucleosome in the absence of rapamycin. **(D)** Left: Metagene analysis of the +Rap/-Rap fold-change ofHir2 ChEC in the Nrdl-AA strain. Profiles were centered around the TATA or TATA-like Binding Site (TBS). The grey box represents the 100bp-centered area in which statistics are generated. Right: Boxplot depicting the +Rap/-Rap fold-change of Hir2 ChEC in the 100bp-TBS centered area. Asterisks indicate significant differences according to a two-tailed Mann-Whitney test. **(E)** Same as in **(D)** with incorporation of newly synthesized H3^HA^ in a 100bp-TBS centered area. **(F)** Snapshot of the MNase-seq profile in the Nrdl-AA-Rap (top) and of the +Rap/-Rap fold­ changes of MNase-seq, Hir2 ChEC and newly synthesized H3^HA^ incorporation in the Nrdl-AA strain at the *VAC7* locus. **(G)** ChIP of H3^HA^ in Nrdl-AA and Nrdl-AA *hir2Δ* strains upon rapamycin treatment. Immunoprecipitated iAS and nAS promoters were normalized to immunoprecipitated *SPT15* ORF after qPCR. Fold change was artificially set to 1 for all the-Rap conditions. n=2, error bars represent the SEM. **(H)** Heatmaps of MNase-seq profile in the Nrdl-AA-Rap and of the +Rap/-Rap fold-changes in Hir2 ChEC and newly synthesized H3^HA^ incorporation in the Nrdl-AA strain at the 601 iAS genes. Graphs are centered on the +1 nucleosome. Each heatmap is divided into two categories of NDR widths: Large NDRs (>150bp) in which a nucleosome can virtually be incorporated, and Small NDRs in which a nucleosome cannot fit. **(I)** Molecular model of the link between Sense/Antisense transcription, HIR binding and H3 incorporation into chromatin. Transcription facilitates stochastic nucleosome loss that can be compensated by incorporation of H3 by the HIR complex. When antisense is induced, nucleosomes are repositioned leading to a stochastic loss and replacement of nucleosomes according to this new phasing.

AMTI occurs through antisense extension over promoters; hence we repeated our analysis focusing on the promoter NDRs. We found that antisense induction leads to a significant higher recruitment of both Hir2 and incorporation of soluble H3 over the TBS for the iAS as compared to the nAS (Figures 3D, 3E and 3F). Moreover, the increased H3^HA^ incorporation at promoters upon antisense extension is Hir2-dependent (Figure 3G). Increased Hir2 binding and H3^HA^ incorporation are not restricted to large NDRs since even <150bp NDRs, which cannot accommodate a nucleosome, present such increases (Figure 3H). Thus, increased incorporation does not reflect the addition of an extra nucleosome into the NDR but rather the replacement by HIR of a former nucleosome that has first been shifted by antisense extension (Figure 3I).

Altogether, our results indicate a positive correlation between coding/non-coding transcription, HIR binding and replication independent H3 incorporation (Figure S2E). However, it is worth noting that increased recruitment of Hir2 and H3^HA^ incorporation at promoters is strictly dependent on increased antisense transcription and not on the SAGA- or TFIID-dependent nature of genes (Figures S2F and S2G).

### SAGA-dependent genes at steady state are enriched in antisense, HIR binding and H3 incorporation at promoter NDRs

Only a small number of iAS genes are SAGA-dependent (Figures 2D and 2E). However, we recently proposed that 20% of the *S. cerevisiae* coding genes are influenced by steady-state antisense transcription into promoters. Thus, some of the SAGA-dependent genes might have a high natural level of antisense that does not further increase upon Nrd1 anchor-away, and are therefore not classified as iAS genes. Thus, we decided to consider the whole set of SAGA-dependent genes in order to assess their natural antisense levels as well as their dependency on the HIR complex for coding gene expression.

We found that SAGA-regulated genes (549 genes) present a significantly higher steady-state level of nascent antisense transcription into promoters as compared to TFIID-dominated genes (3,998 genes) (Figure 4A, left panel), despite showing no difference in coding sense transcription (Figure 4A, right panel). This enrichment of steady-state antisense transcription is well illustrated by analysis of nascent transcription at the *PHO84* (Figure 4B). In agreement with our model of promoter NDR closing by antisense transcription and as already described in the literature, promoters of SAGA-regulated genes show a higher density of nucleosomes over the TBS (Figure 4C). Since we have shown a positive correlation between increase in antisense transcription and increase of HIR binding at promoters (Figure 3), one could expect higher steady-state recruitment of HIR at SAGA-dependent genes as compared to TFIID-regulated genes. Indeed, steady-state Hir2 binding and incorporation of soluble H3^HA^ are significantly higher at SAGA-dependent gene promoters (Figures 4D and 4E).

**Figure 4:**
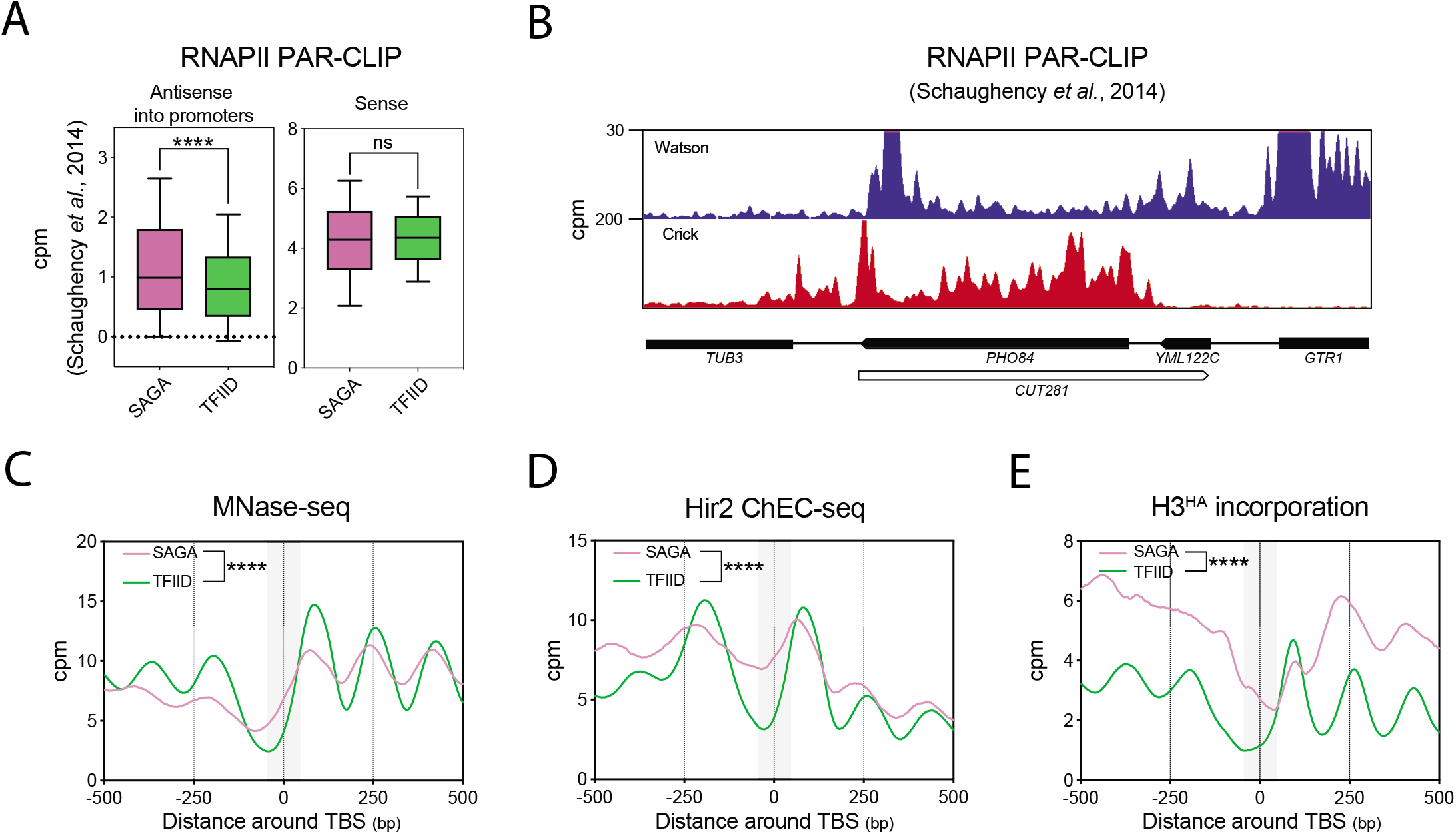
SAGA-dependent genes present high levels of antisense, closed NDRs, high levels of HIR binding and H3 incorporation at promoters. **(A)** Levels of nascent antisense into promoters (−100 to TSS area) and sense transcription (−100 to polyadenylation site) as revealed by RNAPII PAR-CLIP at 549 SAGA-dependent genes and 3,998 TFIID-dependent genes. Data were taken from Schaughency et al., 2014. **(B)** Snapshot depicting nascent transcription revealed by RNAPII PAR-CLIP in sense and antisense orientation at *PH0 84*. **(C)** Metagene plot of nucleosome positioning assessed by MNase-seq in an Nrdl-AA strain in-Rap condition at the 549 SAGA- and 3,998 TFDII-dependent genes. The grey box represents the −50 to +50bp TBS-centered area in which statistics were generated. Asterisks indicate significant differences according to a two-tailed Mann-Whitney test. **(D)** and **(E)** Same as in **(C)** with Hir2 ChEC-seq and H3^HA^ incorporation profiles.

### HIR represses SAGA-dependent genes *via* promoter NDR closing

If HIR binds more SAGA-dependent gene promoters, thereby increasing H3 *de novo* deposition in order to maintain the promoters in a closed state, its deletion might lead to chromatin opening and subsequent gene expression up-regulation. In agreement with such a statement, SAGA-dependent genes are up-regulated in the absence of Hir2 (Figure 5A, left panel). This gene de-repression is not due to an altered antisense production (Figure 5A, right panel) but to a decrease in nucleosome occupancy over the TBS and a subsequent increased PIC recruitment (Figures 5B, 5C and 5D).

**Figure 5:**
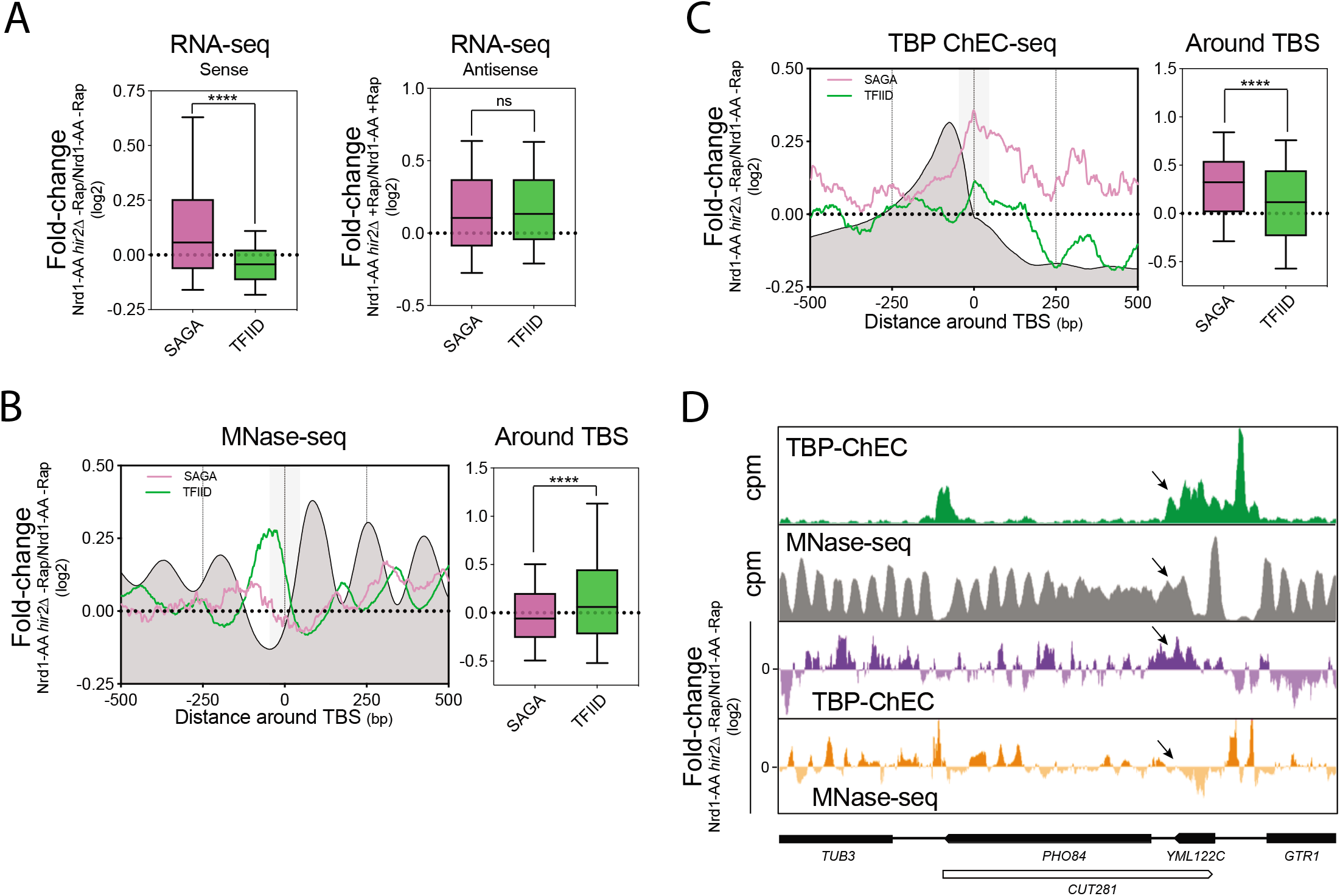
HIR-dependent closing of promoter NDRs represses SAGA-regulated genes. **(A)** Boxplot showing the Nrdl-AA *hir2Δ*/Nrd1-AA fold-change in RNA-seq for the 549 SAGA- and 3,998 TFIID-dependent genes. The whole sense transcription unit is considered for calculation of fold-changes in sense and antisense. Asterisks indicate significant differences according to a two-tailed Mann-Whitney test. **(B)** Left: Metagene analysis of the Nrdl-AA *hir2Δ*/Nrd1-AA fold-change in MNase-seq. The dark grey profile represents the average nucleosome in the absence of rapamycin. The grey box represents the 100bp TBS-centered area in which statistics are generated. Right: Boxplot depicting the Nrdl-AA *hir2Δ*/Nrd1-AA fold-change in MNase-seq in the l00bp TBS-centered area. Asterisks indicate significant differences according to a two-tailed Mann-Whitney test. **(C)** Same as in **(B)** with signal coming from TBP-ChEC. The dark grey represents the TBP­ ChEC signal in the Nrdl-AA strain. **(D)** Snapshot depicting the *PH084* gene for TBP-ChEC and MNase-seq. Arrows indicate the position of the *PH084* TBS.

Altogether, we conclude that HIR binding at promoters of SAGA-regulated genes is enhanced by antisense transcription. This recruitment is important for repression of SAGA-dependent genes by favoring nucleosome incorporation instead of transcription initiation machinery binding.

### The balance between nucleosome incorporation and TF binding at promoters is an important feature to define the SAGA- or TFIID-dependent nature of genes

SAGA-dependent genes show higher levels of antisense into their promoters (Figure 4A). However, some TFIID-dependent genes show equivalent levels of antisense into their promoters without being repressed by HIR (Figures S3A and S3B). Moreover, induced antisense increases HIR binding and H3 incorporation at promoters independently of the gene classes (Figures S2F and S2G). Thus, if higher level of antisense transcription into promoters is a feature of SAGA-dependent genes, an additional feature might help in the definition of the gene class. An earlier model proposed a highest competition between nucleosome and TFs on chromatin at SAGA-dependent genes (23). Thus, we hypothesized that changing the balance between nucleosome incorporation and TF binding might turn a HIR/SAGA-dependent gene into a HIR-independent gene and *vice versa* (Figure 6A).

**Figure 6:**
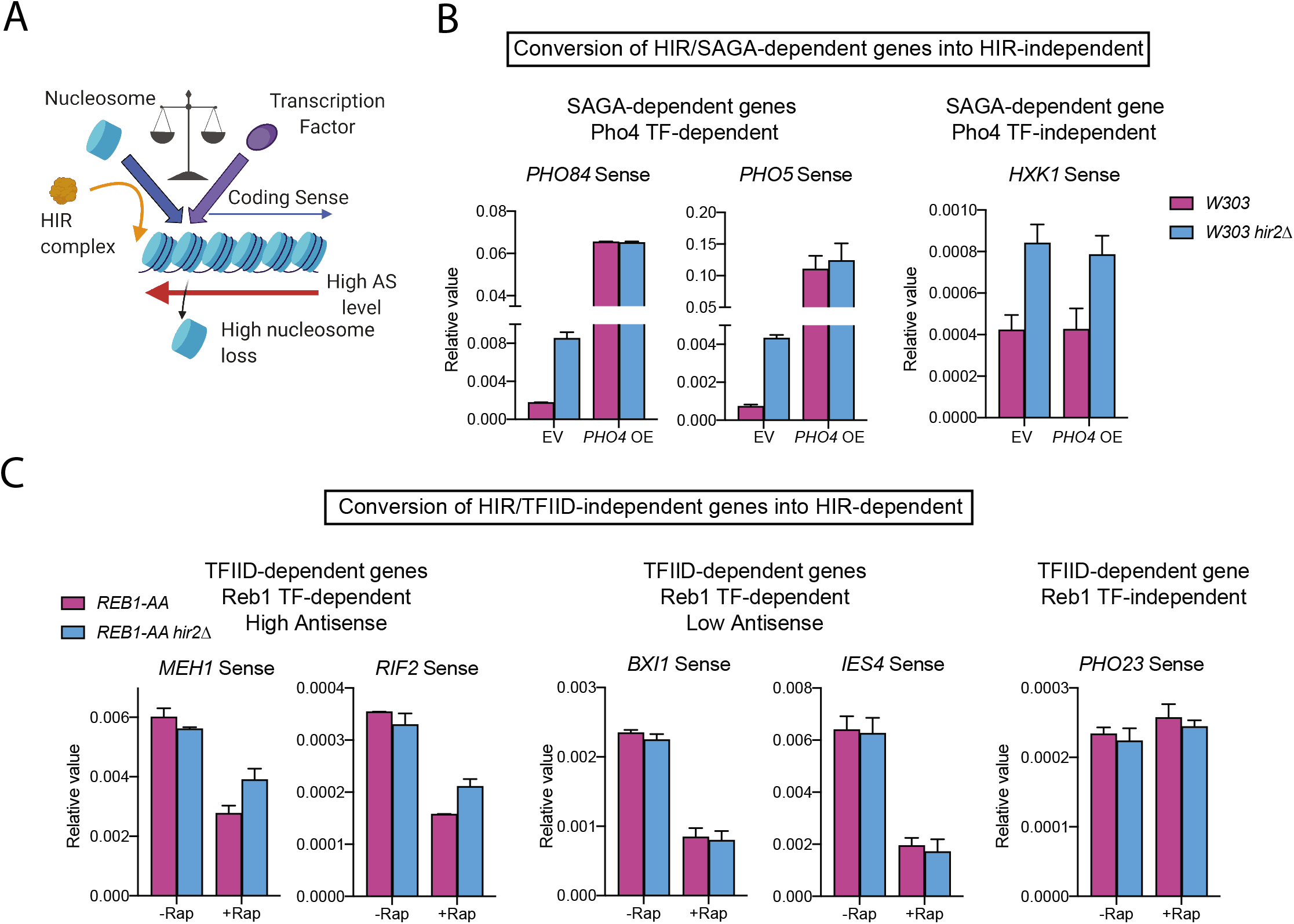
BIR-dependent closing of promoter NDRs represses SAGA-regulated genes. **(A)** Antisense transcription into promoters increases nucleosome loss. We hypothesize that this loss may be compensated either by (H3-H4)_2_ incorporation by HIR (SAGA-dependent genes), or transcription factor binding (TFIID-dependent genes). Thus, the balance between the two mechanisms in competition may be a novel criterion distinguishing the SAGA- or TFIID­ dependent gene classes. **(B)** RT-qPCR analyses of *PH0 84, PH05* and *HXKI* sense expression normalized over *SCR1* expression in the indicated strains. n=2, error bars represent the Standard Error of the Mean (sem). EV: Empty 2-micron plasmid, *PH04* OE: overexpression of *PH04* inserted in a 2-micron plasmid. **(C)** RT-qPCR analyses of *MEHi, RIF2, BXII, IES4* and *PH023* sense expression normalized over *SCR1* expression in the indicated strains. Reb1 depletion from the nucleus upon rapamycin addition was induced for 30min. n=2, error bars represent the SEM.

We first overexpressed the Pho4 TF, which regulates *PHO84* and *PHO5* expression, by introducing an extra copy of its gene on a 2-micron plasmid. While *PHO84* and *PHO5* are repressed by HIR in the presence of an empty vector, they become overexpressed and independent of HIR upon Pho4 overexpression (Figure 6B). As expected, Pho4 up-regulation does not affect SAGA-dependent genes, such as *HXK1*, that are not targeted by this transcription factor. Hence, the overexpression of the TF is able to convert a HIR/SAGA-dependent gene into HIR-independent.

We then selected TFIID-dependent genes with high and low steady state levels of antisense into promoters and which are regulated by the Reb1 TF (44) (Figure S3C). For this experiment, we induced Reb1 depletion from the nucleus using the anchor-away system. On one hand, *MEH1* and *RIF2* which have high steady-state levels of antisense into promoters are down-regulated following Reb1 depletion (Figure 6C). However, upon deletion of *HIR2*, this down-regulation is lesser than in the presence of the histone chaperone. This is not the case for the *BXI* and *IES4* genes, which have low levels of antisense into promoters, and are therefore not dependent on the HIR complex for gene regulation. As a control, we used *PHO23* as a Reb1-independent gene. Thus, by rarefying the TF Reb1, genes with high antisense into their promoters become more prone to gene expression upon *HIR2* deletion, hence turning a HIR-independent into a HIR-dependent gene.

## Discussion

Starting with a genetic screen, our work has led to establishing a strong functional link between antisense transcription, the binding of the HIR histone chaperone at promoters and SAGA-/TFIID-dependent gene regulation. First, we have shown that antisense induction leads to nucleosome repositioning all along the coding region, but also at promoters, and the subsequent displacement triggered by HIR binding and *de novo* histone deposition (Figure 3). Second, SAGA-dependent genes tend to present higher levels of natural antisense into promoters, a narrower NDR, as well as higher HIR binding and *de novo* histone deposition (Figure 4). Third, the absence of HIR opens promoter NDRs at SAGA-dependent genes leading to their up-regulation (Figure 5). At last, we have shown that the balance between *de novo* histone deposition and transcription factor binding at promoters is an important feature defining the SAGA- and TFIID-dependent nature of genes (Figure 6).

Altogether, we propose that at SAGA-dependent genes, natural antisense transcription into promoters tends to close promoter NDRs with nucleosomes masking the TBS and TF-binding sites (Figure 7). When nucleosomes are randomly lost in promoters upon antisense transcription elongation, the chromatin has the possibility to open before being filled by a *de novo* deposited nucleosome. In the absence of HIR, promoters stay open allowing access to TFs and the PIC. In the case of TFIID-dependent genes, loss of nucleosomes at promoters is rapidly filled by TF binding even in the presence of high level of antisense transciption, making this class insensitive to the absence of HIR.

**Figure 7:**
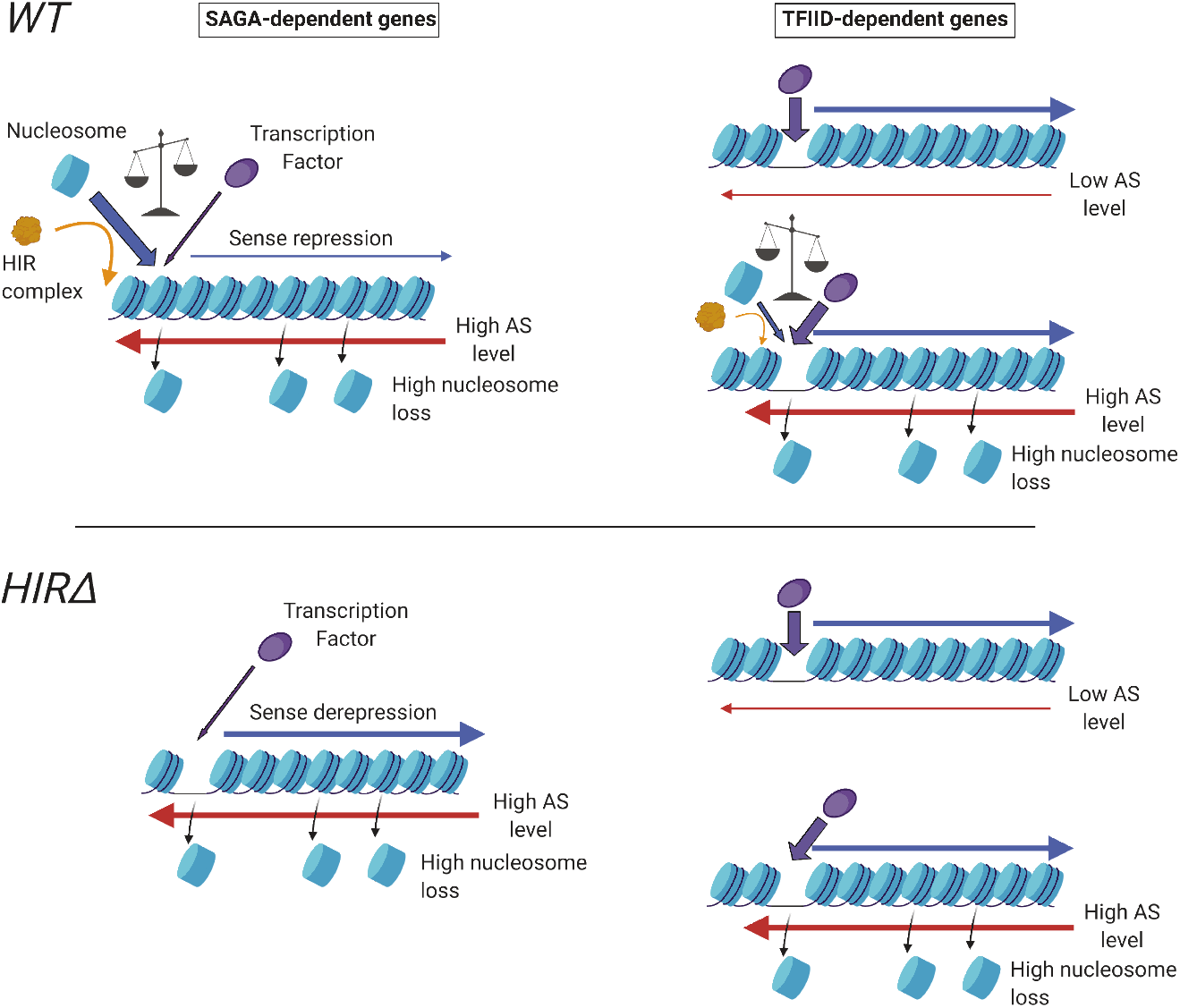
Antisense-mediated transcription interference of SAGA-dependent genes involves the HIR histone chaperone. see discussion

SAGA-dependent genes have been extensively studied and associated with several features. SAGA-dependent promoters present (1) a more closed chromatin structure (23,28–30), (2) nucleosomes undergoing a higher frequency of exchange (26) and (3) a higher responsiveness to the presence of transcription factors (28). These properties can be revisited in the light of our study.

Our data show that SAGA-dependent genes are associated with high levels of steady-state antisense transcription (Figure 4). Antisense transcription into promoters induces a re-phasing of the nucleosomes along the antisense transcription unit (Figure 3) (16). Hence, NDR-flanking nucleosomes are re-positioned, thereby masking important sequences for sense transcription initiation. We therefore propose that steady-state antisense transcription participates in the closing of the chromatin at SAGA-dependent promoters. Moreover, histone exchange is known to be a direct consequence of transcription elongation (26,54), and our model strongly suggests that antisense transcription elongation into SAGA-dependent promoters favors such histone dynamics. At last, the work of the Holstege laboratory has shown that the depletion of Hsf1, a TF targeting genes in the two classes, has a stronger down-regulation effect on the expression of the SAGA-dependent genes (28). Confronting this result to our model, we would propose that SAGA-dependent genes are probably more prone to down-regulation upon Hsf1 depletion because of a competition with the HIR-dependent deposition of nucleosomes into their promoters favored by antisense transcription.

Antisense enrichment in promoters might also contribute to the noisiness of SAGA-dependent transcription (55). Genes with high levels of antisense into their promoters are known to present a higher expression variability (56). Moreover, we have shown that steady-state antisense-enriched genes present a fuzzier organization of nucleosomes. Considering that sense and antisense transcription are mutually exclusive at the single-cell level (50), this fuzziness might represent the mean in the population of the sense- and antisense-dominated chromatin organization for a given gene. Thus, a SAGA-dependent gene might switch back and forth from a sense to antisense dominated organization contributing to the expression heterogeneity in the cell population.

In this study, we did not take into consideration the correlation between the presence of a perfect TATA-box motif and the SAGA-dependent expression of a gene (25,27). Although we observe an enrichment of steady-state antisense levels into promoters of TATA-containing as compared to TATA-less genes, it is less pronounced than the difference between SAGA and TFIID-dependent genes (Figure S4A). Thus, antisense transcription into promoters appears as being more strongly correlated with the coactivator choice than with the TBP turnover at promoters (57,58).

A recent publication has revisited the concept of SAGA- and TFIID-dependent genes with the help of conditional mutants instead of gene deletions and analysis of nascent transcription instead of RNA levels in order to reveal more direct targets of these two regulatory pathways (25). This has led to the new definition of Coactivator-Redundant (CR) genes that are related to SAGA-dependent genes. We observe that these newly defined CR genes are repressed by the HIR complex similarly to the SAGA-dependent genes (Figure S4B).

The rescue by the deletion of the HIR complex appears as relatively weak in our initial genetic screen as compared to the deletion of *RPD3* (Figure 1B). Accordingly, the up-regulation of SAGA-dependent genes upon *HIR2* deletion, while significant, can be considered as low (+15% in mean). It is nevertheless quite consistent with the molecular model we propose. In order to be up-regulated, a SAGA-dependent gene has to lose a nucleosome masking the TF-binding site and the TBS. While the probability of this event is increased by high levels of antisense transcription elongation into promoters, it stays low since the recycling of already incorporated histones by essential histone chaperones is an efficient mechanism (36,59). Thus, such an event is likely to happen only in a minority of cells in the population, when essential histone chaperones are failing, thereby explaining the weak increase in gene expression upon HIR deletion.

Replication-independent histone deposition by HIR occurs within a genetic pathway requiring another upstream histone chaperone, Asf1 (60). Asf1 has been involved in nucleosome turnover during transcription elongation (61). Therefore, loss of Asf1 might as well lead to the up-regulation of SAGA-dependent genes. This is what is observed when using the data from (45), further strengthening our model (Figure S4C).

Interestingly, the HIR complex has also been uncovered in a genetic screen designed to dissect the mechanism of transcription interference when genes are in a *in tandem* configuration (62). In this study, deletion of the HIR complex is rescuing the expression of the downstream gene, *HIS3*. Based on our work, we propose that transcription of the upstream gene into the *HIS3* promoter induces nucleosome loss, resulting in a novel NDR that is not filled by *de novo* nucleosome deposition when HIR is absent. Consequently, the expression of *HIS3*, which is itself a SAGA-dependent gene, increases. It is worth to notice that the HIR complex is also involved in the repression of divergent non-coding transcription (63).

The HIR complex is a major negative regulator of histone expression through recruitment of the Hpc2 subunit to the NEGative regulatory element (NEG) (64,65). We cannot completely rule out that our results are indirect effects due to histone overexpression. However, this possibility is unlikely since the upregulation of SAGA-dependent genes in the absence of HIR is generated by a deficit and not an excess of nucleosomes at promoters.

The typical chromatin organization of SAGA- and TFIID-dependent promoters appears as conserved in human cells and related to the transcriptional plasticity, like in yeast (23). Moreover, the HIR histone chaperone is conserved in metazoans where it is known as the HIRA complex (7). HIRA is involved in the incorporation of the histone variant H3.3 into chromatin in a DNA replication-independent manner. In HeLa cells, the histone chaperone is enriched just upstream of the TSS of highly expressed genes and to a lesser extent in gene bodies with a positive correlation with levels of transcription (66). At last, antisense non-coding transcription into human promoters correlates with more closed promoter NDRs as observed in *S. cerevisiae* (14). Thus, many features of this study are conserved in higher eukaryotes. Altogether, these observations may generate a framework worth of consideration regarding human gene regulation.

## Author Contributions

Conceptualization: J.S., N.B., F.S. Investigation: J.S., N.B, A.M.P., A.M., Z.B. Bioinformatics analyses: J.S., N.B. Writing – Original Draft: J.S. Writing – Review and Editing: J.S., F.S Supervision: F.S. Funding acquisition: F.S.

## Acknowledgements

We thank Guillaume Canal, Michel Strubin and all members of the Stutz laboratory for critical reading of the manuscript, comments and discussions. This work was supported by funds from the Swiss National Science Foundation (grants 31003A 153331 and 31003A_182344 to F.S.) and the Canton of Geneva.

