## Supplementary Material for "Antisense-mediated repression of SAGA-dependent genes involves the HIR histone chaperone"

### Figure S1- related to Figure 2

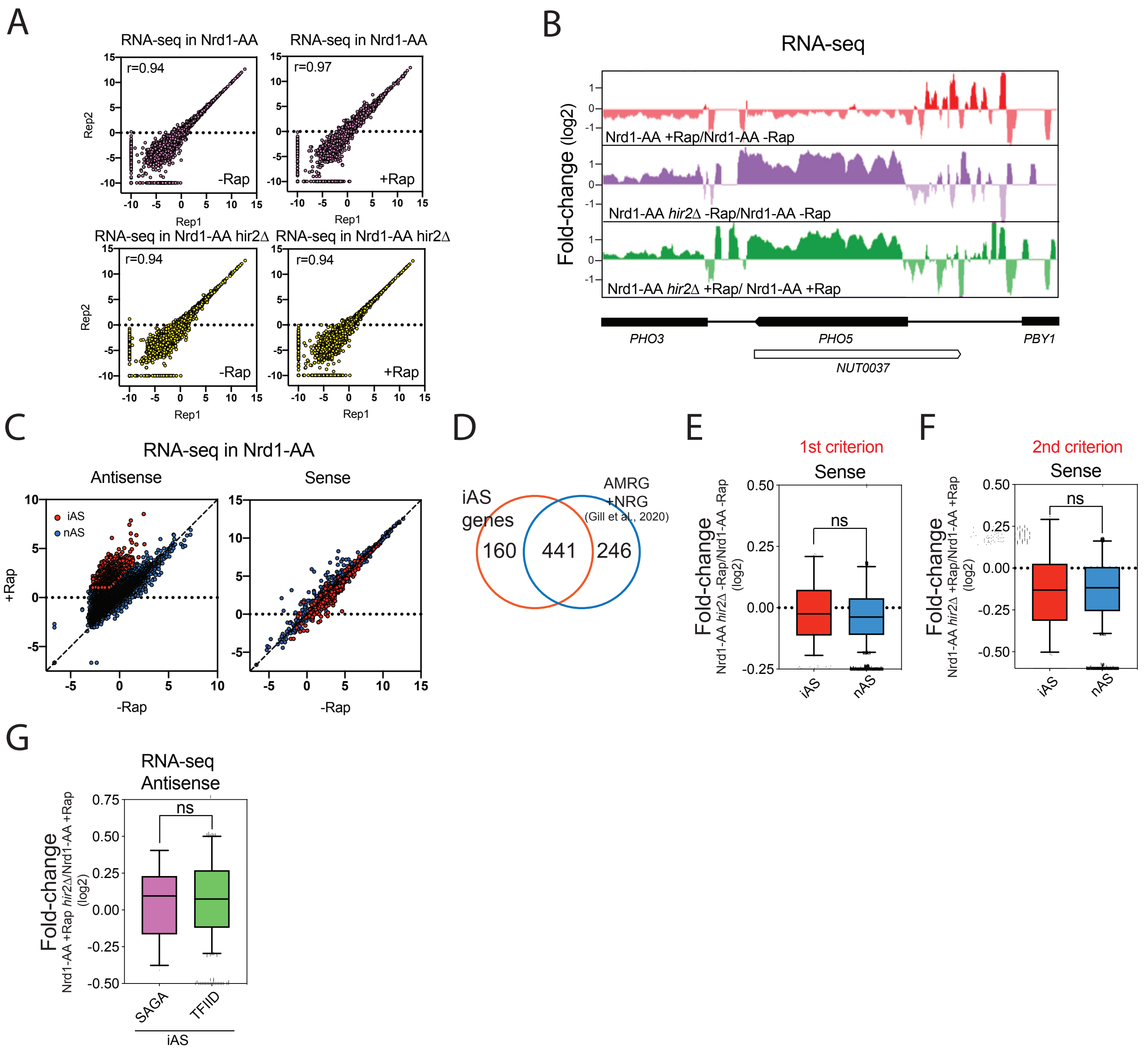


**Figure S1: related to Figure 2**

**(A)** Scatter-plot of correlations between RNA-seq replicates. The Spearman correlation coefficient is indicated.
**(B)** Snapshot of the *PHO5* locus.
**(C)** Scatter-plot of densities for iAS and nAS in Antisense and Sense orientations. Sense RNAs (TSS to polyA) were considered as one bin giving one value. Antisense RNAs correspond to the same measurement on the other strand.
**(D)** Overlap between iAS genes and genes showing increase in Antisense upon Nrd1-AA in our previous study (16).
**(E)** Boxplot showing the fold-change according to the 1st criterion for the iAS and nAS genes.
**(F)** Same as in **(E)**, for the 2nd criterion.
**(G)** Boxplot showing the fold-change (+Rap/-Rap) in Antisense for the iAS SAGA- and TFIID-dependent genes.

### Figure S2- related to Figure 3

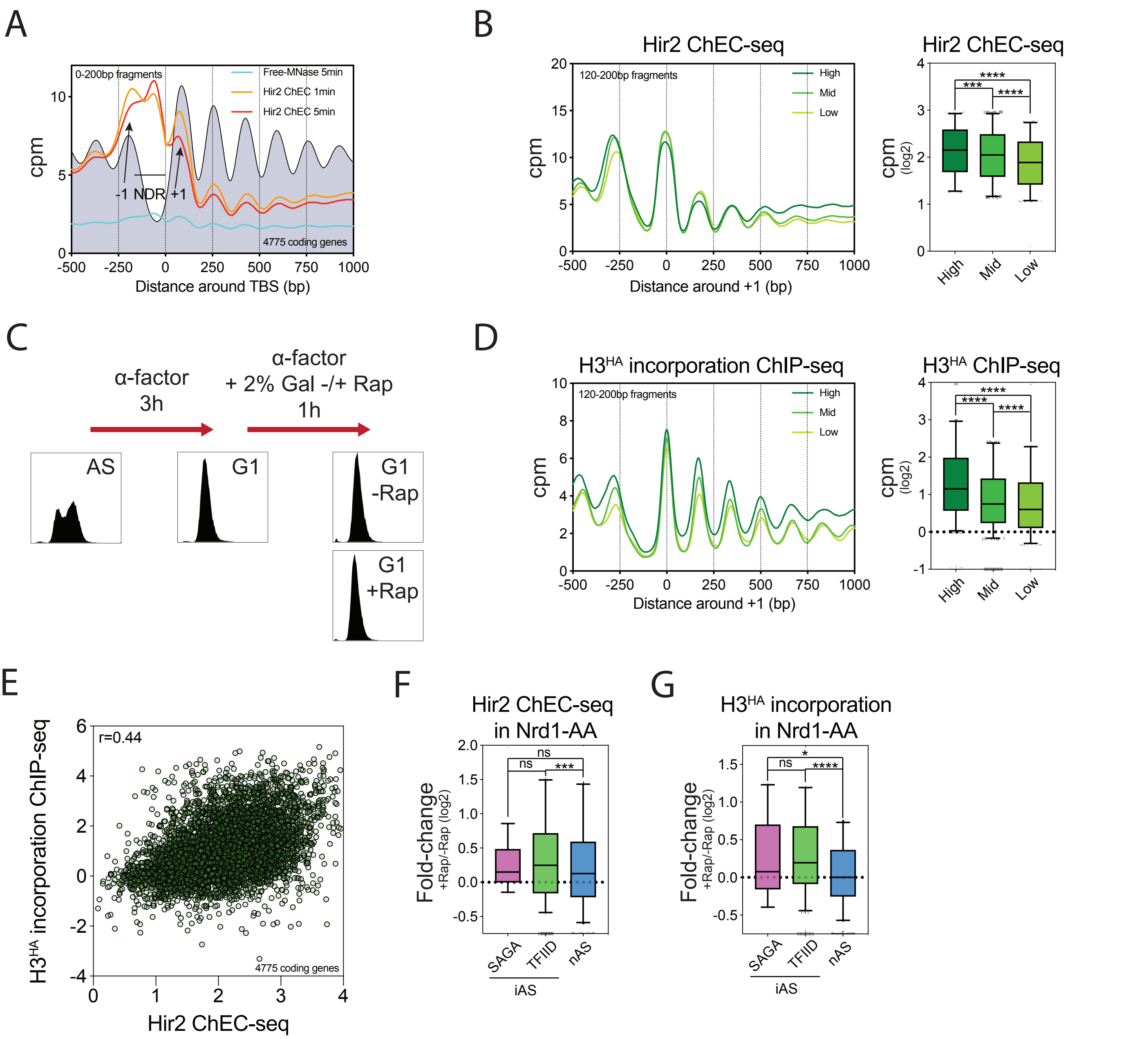


**Figure S2: related to Figure 3**

**(A)** Mapping of the Free-MNase and Hir2 ChEC-seq 0-200bp fragments with respect to the TBS of all coding genes (4,775 genes). The dark grey profile depicts the position of nucleosomes as obtained by MNAse-seq.
**(B)** Left panel: Hir2 ChEC-seq 120-200bp fragments according to +1 nucleosome for Highly (1,592 genes), Midly (1,592 genes) and Lowly transcribed genes (1,591 genes). Right panel: Boxplot of Hir2 ChEC-seq signal for Highly, Midly and Lowly transcribed genes considering the whole gene as one bin.
**(C)** Experimental scheme of the strain culture for the replication-independent $H3^{HA}$ incorporation. Corresponding flow cytometry profiles are indicated.
**(D)** Same as in **(B)**, with the $H3^{HA}$ incorporation signal.
**(E)** Scatter-plot showing the correlation between Hir2 ChEC-seq levels and replication-independent $H3^{HA}$ deposition (4,775 genes). The Spearman correlation coefficient is indicated.
**(F)** Boxplot depicting the +Rap/-Rap fold-change of Hir2 ChEC-seq in the 100bp-TBS centered area making the distinction between iAS SAGA- and TFIID-dependent genes.
**(G)** Same as in **(F)** for the $H3^{HA}$ incorporation signal.

### Figure S3- related to Figure 6

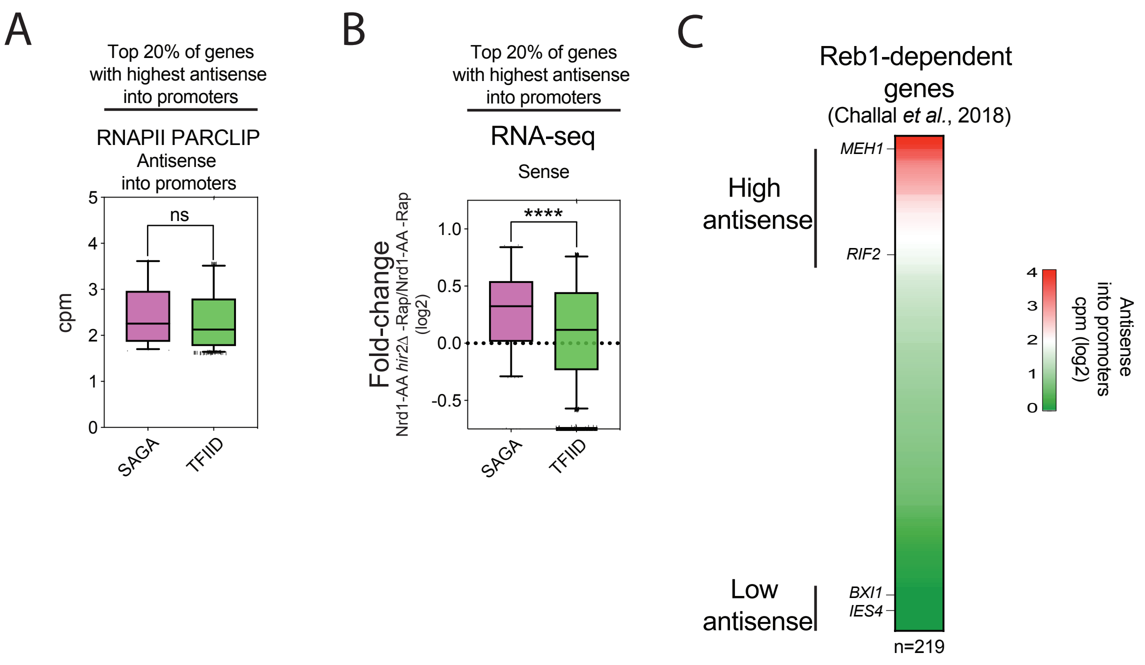


**Figure S3: related to Figure 6**

**(A)** Levels of nascent Antisense into promoters (-100 to TSS area) at SAGA- and TFIID-dependent genes for the quintile of genes showing the highest Antisense level into promoters at steady state. A total of 896 genes were examined of which 168 SAGA-dependent and 728 TFIID-dependent. Data were taken from (69).
**(B)** Same as in **(A)**, but considering the Nrd1-AA *hir2Δ* -Rap/Nrd1-AA -Rap RNA-seq ratio in Sense orientation.
**(C)** Heatmap depicting the natural Antisense level into promoters for the Reb1-dependent genes. Genes used in Figure 6C are indicated.

Figure S4- related to Discussion

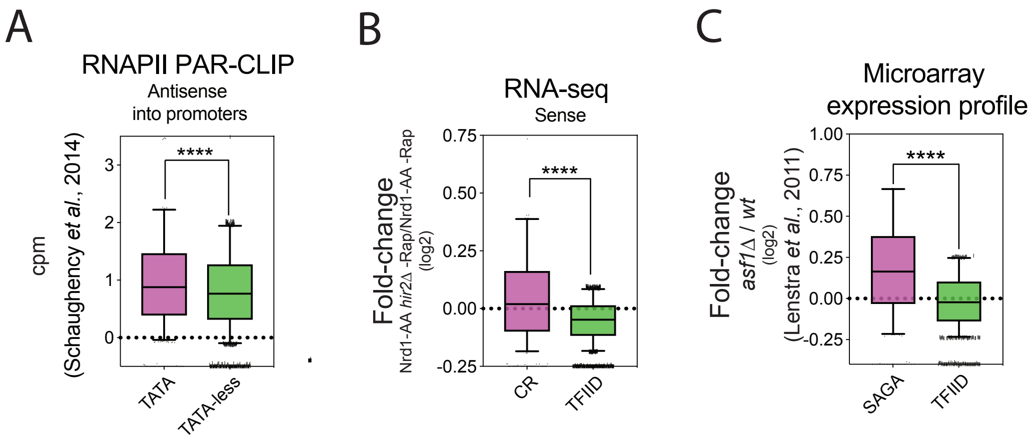


**Figure S4: related to the Discussion**

**(A)** Levels of nascent Antisense into promoters (-100 to TSS area) at 649 TATA and 2985 TATA-less dependent genes.
**(B)** Boxplot depicting the Nrd1-AA *hir2Δ* -Rap/Nrd1-AA -Rap fold-change in Sense of RNA-seq at 523 CR and 3111 TFIID-dependent genes.
**(C)** Boxplot showing the *asf1Δ/wt* fold-change in Sense expression at the 529 SAGA- and 3967 TFIID-dependent genes. Data were retrieved from (45).

#### Table S1

| Systematic name | Standard name |
|---|---|
| YAL028W | FRT2 |
| YAL059W | ECM1 |
| YAL060W | BDH1 |
| YAL067C | SEO1 |
| YAR003W | SWD1 |
| YBL008W | **HIR1** |
| YBL051C | PIN4 |
| YBL080C | PET112 |
| YBR009C | HHF1 |
| YBR025C | OLA1 |
| YBR053C | |
| YBR095C | RXT2 |
| YBR107C | IML3 |
| YBR116C | |
| YBR134W | |
| YBR172C | SMY2 |
| YBR174C | |
| YBR197C | |
| YBR214W | SDS24 |
| YBR215W | **HPC2** |
| YBR225W | |
| YBR238C | |
| YBR264C | YPT10 |
| YCL006C | |
| YCL026C | |
| YCL032W | STE50 |
| YCL037C | SRO9 |
| YCL042W | |
| YCL045C | EMC1 |
| YCL046W | |
| YCL057W | PRD1 |
| YCR016W | |
| YCR022C | |
| YCR037C | PHO87 |
| YCR077C | PAT1 |
| YCR099C | |
| YEL029C | BUD16 |
| YEL031W | SPF1 |
| YER016W | BIM1 |
| YER048C | CAJ1 |
| YER079W | |
| YER089C | PTC2 |
| YER123W | YCK3 |
| YHL029C | OCA5 |

| | |
|---|---|
| YHL035C | VMR1 |
| YHL041W | |
| YHR151C | MTC6 |
| YHR152W | SPO12 |
| YHR179W | OYE2 |
| YHR185C | PFS1 |
| YIL024C | |
| YIL030C | SSM4 |
| YIL040W | APQ12 |
| YIR037W | HYR1 |
| YKL026C | GPX1 |
| YKL069W | |
| YKL081W | TEF4 |
| YKL093W | MBR1 |
| YKL098W | MTC2 |
| YKL149C | DBR1 |
| YKL151C | NNR2 |
| YKR028W | SAP190 |
| YFL063W | |
| YFL052W | ZNF1 |
| YFL033C | RIM15 |
| YFL032W | |
| YFL007W | BLM10 |
| YFL004W | VTC2 |
| YFR040W | SAP155 |
| YDL048C | STP4 |
| YDL124W | |
| YDL167C | NRP1 |
| YDL187C | |
| YDL198C | GGC1 |
| YDL200C | MGT1 |
| YDL229W | SSB1 |
| YDR006C | SOK1 |
| YDR026C | NSI1 |
| YDR048C | |
| YDR057W | YOS9 |
| YDR095C | |
| YDR108W | TRS85 |
| YDR138W | HPR1 |
| YDR210W | |
| YDR214W | AHA1 |
| YDR218C | SPR28 |
| YDR249C | |
| YDR266C | HEL2 |
| YDR289C | RTT103 |
| YDR290W | |

| | |
|---|---|
| YDR318W | MCM21 |
| YDR352W | YPQ2 |
| YDR383C | NKP1 |
| YDR385W | EFT2 |
| YDR415C | |
| YDR424C | DYN2 |
| YDR428C | BNA7 |
| YDR436W | PPZ2 |
| YDR469W | SDC1 |
| YDR475C | JIP4 |
| YDR479C | PEX29 |
| YDR480W | DIG2 |
| YDR494W | RSM28 |
| YDR519W | FPR2 |
| YGL013C | PDR1 |
| YGL131C | SNT2 |
| YGL218W | |
| YGL222C | EDC1 |
| YGL223C | COG1 |
| YGL224C | SDT1 |
| YGL235W | |
| YGR003W | CUL3 |
| YGR004W | PEX31 |
| YGR016W | |
| YGR018C | |
| YGR054W | |
| YGR155W | CYS4 |
| YGR162W | TIF4631 |
| YGR176W | |
| YGR182C | |
| YGR206W | MVB12 |
| YGR250C | RIE1 |
| YGR275W | RTT102 |
| YJL017W | |
| YJL047C | RTT101 |
| YJL068C | |
| YJL198W | PHO90 |
| YJR033C | RAV1 |
| YJR140C | **HIR3** |
| YJR144W | MGM101 |
| YLR034C | SMF3 |
| YLR049C | MLO50 |
| YLR118C | TML25 |
| YLR177W | |
| YLR199C | PBA1 |
| YLR214W | FRE1 |

| | |
|---|---|
| YLR219W | MSC3 |
| YLR283W | PUT7 |
| YLR286C | CTS1 |
| YLR287C | |
| YLR296W | |
| YLR374C | |
| YLR375W | STP3 |
| YLR414C | PUN1 |
| YLR427W | MAG2 |
| YLR436C | ECM30 |
| YLR455W | PDP3 |
| YML054C | CYB2 |
| YML075C | HMG1 |
| YML080W | DUS1 |
| YML086C | ALO1 |
| YML129C | COX14 |
| YMR089C | YTA12 |
| YMR204C | INP1 |
| YMR274C | RCE1 |
| YMR312W | ELP6 |
| YNL040W | |
| YNL053W | MSG5 |
| YNL054W | VAC7 |
| YNL086W | SNN1 |
| YNL097C | PHO23 |
| YNL098C | RAS2 |
| YNL104C | LEU4 |
| YNL141W | AAH1 |
| YNL153C | GIM3 |
| YNL193W | |
| YNR033W | ABZ1 |
| YNR045W | PET494 |
| YOL061W | PRS5 |
| YOL063C | CRT10 |
| YOL088C | MPD2 |
| YOL125W | TRM13 |
| YOR038C | **HIR2** |
| YOR184W | SER1 |
| YOR311C | DGK1 |
| YPL055C | LGE1 |
| YPL066W | RGL1 |
| YPL070W | MUK1 |
| YPL071C | |
| YPL073C | |
| YPL074W | YTA6 |
| YPL077C | |

| | |
|---|---|
| YPL109C | MCO76 |
| YPL110C | GDE1 |
| YPL123C | RNY1 |
| YPL127C | HHO1 |
| YPL139C | UME1 |
| YPL184C | MRN1 |
| YPR043W | RPL43A |
| YPR046W | MCM16 |
| YPR050C | |
| YPR062W | FCY1 |
| YPR070W | MED1 |
| YPR084W | |
| YPR117W | |
| YPR127W | |
| YPR171W | BSP1 |
| YPR179C | HDA3 |

### Table S2- Strain and oligos

| Strains | Genotype | Figures |
|---|---|---|
| WT (BY4742) | MAT α, his3Δ1, leu2Δ0, lys2Δ0, ura3Δ0 | 1A |
| pho84△::HIS3 (FSY4634) | MAT α, pho84Δ::HIS3, leu2Δ0, met15Δ0, ura3Δ0 | 1A, 1B |
| pho84△::HIS3 rrp6△ (FSY4635) | MAT α, pho84Δ::HIS3, rrp6Δ::LEU2, met15Δ0, ura3Δ0 | 1A, 1B |
| pho84△::HIS3 rrp6△ rpd3△ (FSY7764) | MAT α, pho84Δ::HIS3, rrp6Δ::LEU2, rpd3Δ::KanMX6, met15Δ0, ura3Δ0 | 1B |
| pho84△::HIS3 rrp6△ hir1△ (FSY7759) | MAT α, pho84Δ::HIS3, rrp6Δ::LEU2, hir1Δ::KanMX6, met15Δ0, ura3Δ0 | 1B |
| pho84△::HIS3 rrp6△ hir2△ (FSY7761) | MAT α, pho84Δ::HIS3, rrp6Δ::LEU2, hir2Δ::KanMX6, met15Δ0, ura3Δ0 | 1B |
| pho84△::HIS3 rrp6△ hir3△ (FSY7762) | MAT α, pho84Δ::HIS3, rrp6Δ::LEU2, hir3Δ::KanMX6, met15Δ0, ura3Δ0 | 1B |
| pho84△::HIS3 rrp6△ hpc2△ (FSY7763) | MAT α, pho84Δ::HIS3, rrp6Δ::LEU2, hpc2Δ::KanMX6, met15Δ0, ura3Δ0 | 1B |
| pho84△::HIS3-NatMX6, rrp6△ | MAT α, can1Δ::STE2pr-URA3, lyp1Δ, ura3Δ0, met15Δ0, pho84Δ::HIS3-NatMX6, rrp6Δ::LEU2 | 1C |
| W303 RRP6 (FSY4976) | MAT a, leu2-3,112, trp1-1, can1-100, ura3-1, ade2-1, his3-11,15 | 1E, 1F |
| hir1△ RRP6 (FSY5331) | MAT a, hir1△::LEU2, trp1-1, can1-100, ura3-1, ade2-1, his3-11,15 | 1E, 1F |
| hir2△ RRP6 (FSY5333) | MAT a, hir2△::LEU2, trp1-1, can1-100, ura3-1, ade2-1, his3-11,15 | 1E, 1F |
| W303 rrp6△ (FSY3117) | MAT a, rrp6::KanMX6, leu2-3,112, trp1-1, can1-100, ura3-1, ade2-1, his3-11,15 | 1E, 1F |
| hir1△ rrp6△ (FSY5332) | MAT a, hir1△::LEU2, rrp6::KanMX6, trp1-1, can1-100, ura3-1, ade2-1, his3-11,15 | 1E, 1F |
| hir2△ rrp6△ (FSY5334) | MAT a, hir2△::LEU2, rrp6::KanMX6, trp1-1, can1-100, ura3-1, ade2-1, his3-11,15 | 1E, 1F |

| | | |
|---|---|---|
| Nrd1-AA (FSY5329) | MAT a, tor1-1, fpr1::NAT, RPL13A-2×FKBP12::TRP1, NRD1-FRB::KanMX6, bar1::LEU2, his3-11,15, ura3-1, ade2-1 | 2, 5A, 5B, 5D, S1 |
| Nrd1-AA hir2△ (FSY8647) | MAT a, tor1-1, fpr1::NAT, RPL13A-2×FKBP12::TRP1, NRD1-FRB::KanMX6, bar1::LEU2, hir2△::HIS3, ura3-1, ade2-1 | 2, 5A, 5B, 5D, S1 |
| Nrd1-AA Free-MNase (FSY8967) | MAT a, tor1-1, fpr1::NAT, RPL13A-2×FKBP12::TRP1, NRD1-FRB::KanMX6, bar1::LEU2, pREB1-3XFLAG-MNase-URA3, his3-11,15, ade2-1 | S2A |
| Nrd1-AA HIR2-MNase (FSY8968) | MAT a, tor1-1, fpr1::NAT, RPL13A-2×FKBP12::TRP1, NRD1-FRB::KanMX6, bar1::LEU2, HIR2-3XFLAG-MNase-HphMX6, his3-11,15, ura3-1, ade2-1 | 2B, 2D, 2F, 2H, 3D, S2A, S2B, S2E, S2F |
| Nrd1-AA pGAL-H3-3XHA-URA3 | MAT a, tor1-1, fpr1::NAT, RPL13A-2×FKBP12::TRP1, NRD1-FRB::KanMX6, bar1::LEU2, his3-11,15, ade2-1 pRS316-pGAL1-H3-3XHA-URA3 | 3C, 3E, 3F, 3G, 3H, 4E, S2D, S2G |
| Nrd1-AA hir2△ pGAL-H3-3XHA-URA3 | MAT a, tor1-1, fpr1::NAT, RPL13A-2×FKBP12::TRP1, NRD1-FRB::KanMX6, bar1::LEU2, hir2△::HIS3, ade2-1 pRS316-pGAL1-H3-3XHA-URA3 | 3G |
| Nrd1-AA TBP-MNase (FSY8645) | MAT a, tor1-1, fpr1::NAT, RPL13A-2×FKBP12::TRP1, NRD1-FRB::KanMX6, bar1::LEU2, SPT15-3XFLAG-MNase-HphMX6, his3-11,15, ura3-1, ade2-1 | 5C |

| | | |
|---|---|---|
| Nrd1-AA TBP-MNase hir2△ (FSY8646) | MAT a, tor1-1, fpr1::NAT, RPL13A-2×FKBP12::TRP1, NRD1-FRB::KanMX6, bar1::LEU2, SPT15-3XFLAG-MNase-HphMX6, hir2△::HIS3, ura3-1, ade2-1 | 5C |
| W303 pRS426-Empty (FSY9049) | MAT a, leu2-3,112, trp1-1, can1-100, ura3-1, ade2-1, his3-11,15 + 2micron pRS426-URA3 | 6B |
| W303 pRS426-PHO4 (FSY9051) | MAT a, leu2-3,112, trp1-1, can1-100, ura3-1, ade2-1, his3-11,15 + 2micron pRS426-PHO4-URA3 | 6B |
| W303 hir2△ pRS426-Empty (FSY9047) | MAT a, hir2△::LEU2, trp1-1, can1-100, ura3-1, ade2-1, his3-11,15 + 2micron pRS426-URA3 | 6B |
| W303 hir2△ pRS426-PHO4 (FSY9048) | MAT a, hir2△::LEU2, trp1-1, can1-100, ura3-1, ade2-1, his3-11,15 + 2micron pRS426-PHO4-URA3 | 6B |
| Reb1-AA (FSY8558) | MATa tor1-1 fpr1::loxP-LEU2-loxP RPL13A-2×FKBP12::loxP, REB1-FRB::KanMX6, bar1::URA3, ade 2-1 trp1-1 can1-100, his3-11,15 | 6C |
| Reb1-AA hir2△ (FSY9046) | MATa tor1-1 fpr1::loxP-LEU2-loxP RPL13A-2×FKBP12::loxP, REB1-FRB::KanMX6, hir2△::HIS3, ade 2-1 trp1-1 can1-100 | 6C |

| Oligos | Sequence |
|---|---|
| PHO84_Fwd | CCGTCAATAAAGATACTATTCATGTTGCTG |
| PHO84_Rev | AAAATCATTCAAATGGTTGTGGAAGGC |
| PHO5_Fwd | CTTGGGACTACGATGCCAA |
| PHO5_Rev | ACTTCAAATGCACACCACGA |
| SCR1_Fwd | AACCGTCTTTCCTCCGTCGTAA |

| | |
|---|---|
| SCR1_Rev | CTACCTTGCCGCACCAG ACA |
| VAC7_Fwd | AACATCGGCTCGAATAC ACC |
| VAC7_Rev | AGCCCATCATTTGCTACT GG |
| PHO23_Fwd | ACGAGAGCCAAGACCA CACT |
| PHO23_Rev | GTTTCCGACGCTGCTAA CTC |
| EPL1_Fwd | TACGGTCGAACAAAGG GAAG |
| EPL1_Rev | AACAGTTGCTGCTGTTG CTG |
| RAM2_Fwd | TTTCAACATGAGCTCGC CTA |
| RAM2_Rev | TGCGGAACCAATTGAAT CTT |
| SPT15_Fwd | GCCAACAACAAGCTGA ATTCAAGC |
| SPT15_Rev | CCACCGTATTCGCGGGC ATTTG |
| HXK1_Fwd | ATGGACCAAGGGTTTCG ATATTCC |
| HXK1_Rev | ATCTTAGTCTCTGGGTCA GTGTAGT |
| MEH1_Fwd | ACAAATGCGGTTGAAGG AAC |
| MEH1_Rev | CCTGCACTCCTTTCGTCT TC |
| RIF2_Fwd | GCGAGTGGACCATGTTT TCT |
| RIF2_Rev | CTGAAACTGCACGGTCG ATA |
| BXI1_Fwd | GCGGTTTATGCACAAGG TTT |
| BXI1_Rev | CTCGTAGTCTTCGGGAC GAG |
| IES4_Fwd | AGGATACAAGCCCTCCC TCT |
| IES4_Rev | TAAACTCGCTTCCCCTTT GA |
